# Collablots: Quantification of collagen VI levels and its structural disorganisation in cell cultures from patients with collagen VI-related dystrophies

**DOI:** 10.1101/2024.12.06.627141

**Authors:** Nadia Osegui-Barcenilla, Maria Sendino, Sergio Martín-González, Itziar Gómez-Moro, Ainhoa Benito-Agustino, Noemi Torres-Conde, Andrea López-Martínez, Cecilia Jiménez-Mallebrera, Arístides López-Márquez, Virginia Arechavala-Gomeza

**Author notes:** **Corresponding Author:** Prof. Virginia Arechavala-Gomeza Nucleic Acid Therapeutics for Rare Diseases, Biobizkaia Health Research Institute, Plaza de Cruces S/N, 48903 Barakaldo, Spain.

## Abstract

**Aims:** This study aims to develop a quantitative method for assessing collagen VI expression in cell cultures, which is crucial for the diagnosis and treatment of collagen VI-related dystrophies.

**Methods:** We developed a combined in-cell western (ICW) and on-cell western (OCW) assay, that we have called ‘collablot’ to quantify collagen VI and its organisation in the extracellular matrix of cell cultures from patients and healthy controls. To optimise it, we optimised cell density and the protocols to induce collagen expression in cultures, as well as the cell fixation and permeabilisation methods. This was completed with a thorough selection of collagen antibodies and a collagen hybridising peptide (CHP). We then used collablots to compare cultures from patients and controls and evaluate therapeutic interventions in the cultures.

**Results:** Collablots enabled the quantification of collagen VI expression in both control and patient cells, aligning with immunocytochemistry findings and detecting variations in collagen VI expression following treatment of the cultures. Additionally, CHP analysis revealed a marked increase in collagen network disruption in patients compared to the controls.

**Conclusions:** The collablot assay represents an optimal method for quantifying collagen VI expression and its organisation in culture and assessing the effect of therapies.

**Key Points:**

- Evaluating therapies for collagen VI-related dystrophies (COL6-RD) requires the quantification of collagen VI levels.
- Collablot assays are a novel method for quantifying collagen VI expression and its structural organisation in cell culture.
- Due to the significant role of phenotype heterogeneity in this complex disease, quantifying collagen alone might not be adequate for diagnosing COL6-RD, but the addition of a peptide to quantify collagen disorganisation could help in the characterisation of patient cultures.

## Introduction

Myopathies associated with collagen VI deficiency are caused by one or more pathogenic variants in the *COL6A1*, *COL6A2*, and/or *COL6A3* genes, which encode the three essential α-chains of collagen VI. These mutations result in a spectrum of diseases known as collagen VI-related dystrophies (COL6-RDs), ranging from relatively mild Bethlem muscular dystrophy (BM) to severe Ullrich congenital muscular dystrophy (UCMD). Intermediate phenotypes, referred to as intermediate COL6-RD, represent the clinical spectrum observed between the two extremes previously mentioned^1^.

In muscle, collagen VI functions as a ubiquitous extracellular matrix (ECM) protein within the stroma, constructing a microfibrillar network linked to the basement membrane^2^ It is synthesised not by myocytes, but by the resident primary interstitial muscle fibroblasts in the musculoskeletal system^3^. Inside these fibroblasts, the three distinct chains come together to form trimeric structures, subsequently organised into antiparallel dimers and eventually assembled into tetramers. These tetramers are secreted into the extracellular matrix to form the previously mentioned microfibril network^4^ (Figure 1). Mutations in the genes encoding essential chains disrupt trimer formation, causing intracellular collagen retention and impaired microfibril network formation.

**Figure 1.**
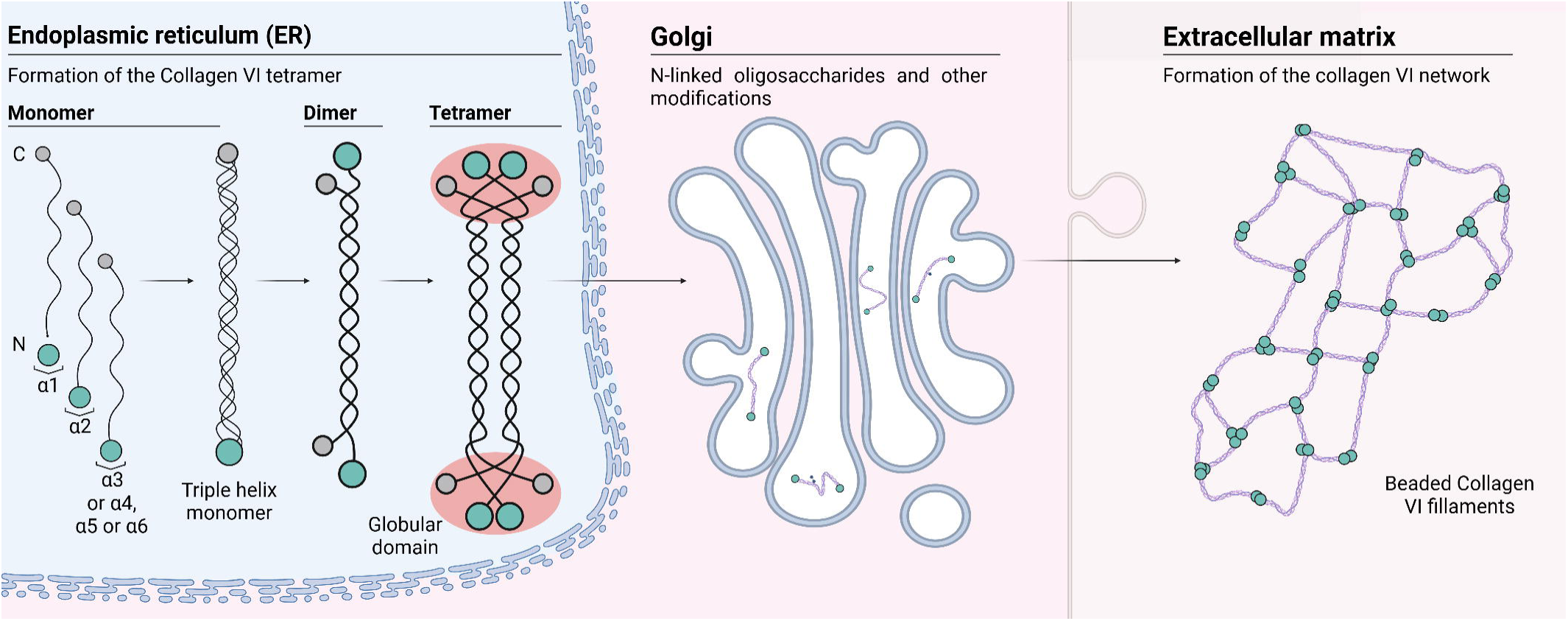
Collagen VI assembly and secretion.

While formal diagnostic criteria for collagen VI-related dystrophies have not yet been established, diagnosis typically involves clinical evaluation, muscle biopsy analysis, and identification of mutations in the relevant genes using molecular genetic testing^1^. However, achieving a genetic diagnosis can be challenging due to the large size of the *COL6A* genes (*COL6A1*: 23,281bp, *COL6A2*: 34,737bp, *COL6A3*: 40,493bp) and their high frequency of polymorphisms. Despite being invasive, muscle biopsies remain the sole method to assess collagen expression, albeit qualitatively.

Several approaches have been proposed to genetically treat collagen VI-related dystrophies, with most strategies focusing on RNA molecules or gene editing techniques^5–8^. However, to assess the efficacy of these putative therapies, there is currently no validated method for quantifying collagen expression in the cell cultures that are routinely used to develop these treatments.

Efforts have been made to develop methods for studying collagen VI expression, but most lack quantitative accuracy. The primary method for assessing this protein in patients’ muscles is direct labelling of muscle biopsies with antibodies against collagen VI; this approach can localise and show a decrease in the expression of collagen VI. However, obtaining muscle biopsies is often challenging due to their invasive nature. In such cases, immunolabeling studies are sometimes performed on skin biopsies. Both methods exhibit high sensitivity and specificity in diagnosing Ullrich muscular dystrophy (close to 100%), with lower specificity for Bethlem myopathy (63%). Therefore, while collagen VI immunostaining is a valuable diagnostic tool, particularly for Ullrich cases, is not a quantitative technique and heavily relies on the expertise of the pathologist analysing the sample^9,10^.

*In vitro* studies can also be conducted using skin-derived fibroblasts. Conventional analysis of collagen VI typically involves immunohistochemistry, a method that can reveal a reduction or abnormal deposition of the extracellular matrix of collagen VI, along with increased intracellular retention observed upon cell permeabilisation. However, this method is both laborious and non-quantitative^11^. Another *in vitro* analysis using fibroblasts is flow cytometry, which provides quantitative data, but requires a large number of cells, often difficult to obtain or requiring prolonged culture periods^12^.

We have prior experience in developing methods to quantify proteins associated with other neuromuscular disorders using in-cell western (ICW) technology^13,14^. An in-cell western is a quantitative immunofluorescence assay performed in microplates allowing for the quantification of proteins directly within cell cultures. Collagen VI is a protein produced intracellularly but excreted and organised in the extracellular matrix. To accurately quantify this protein, we have combined ICWs, which target intracellular proteins thanks to cell permeabilisation, with on-cell westerns (OCW), where samples are not permeabilised, hence staining only the secreted portion of the protein. Additionally, we assessed the integration of a ligand to measure collagen disorganisation within the same system: collagen hybridising peptides (CHPs) demonstrate high binding specificity to denatured collagen chains while exhibiting very low affinity for intact (triple helical) collagen^15^. We have optimised this assay, which we call ‘collablot’, to effectively evaluate variations in protein expression in patient cultures and provide an *in vitro* screening method for potential therapies.

## Material and methods

### Samples

Primary fibroblasts were used to refine and evaluate the protocol. These cells, derived from skin biopsies of healthy controls and COL6-RD patients collected after informed consent (Table 1), were either purchased from commercial suppliers (CTRL FP, catalogue number CC-2511, Lonza Bioscience, Morrisville, NC, USA), obtained from Biogipuzkoa Health Research Institute (Donostia-San Sebastian, Spain) or sourced from the Biobank of the Hospital Sant Joan de Déu (BHISJDI, Barcelona, Spain). The previously generated edited culture pair (COLVI FP0016 and COLVI FP0016edit), derived from a COL6-RD patient with an intermediate phenotype, was provided by Drs Jiménez-Mallebrera and López-Márquez^16^.

**Table 1.** Skin-derived fibroblast cultures used in this study.

| Sample culture | Mutation | Severity | Culture | Doubling time (days) | Origin |
| --- | --- | --- | --- | --- | --- |
| **Controls** |  |  |  |  |  |
| CTRL FP0821 | None | None | Primary | 2.1 | Biogipuzkoa HRI |
| CTRL FP | None | None | Primary | 2.3 | Lonza |
| CTRL FP0010 | None | None | Primary | 1.9 | Sant Joan de Déu |
| CTRL FP0017 | None | None | Primary | 3.1 | Sant Joan de Déu |
| **Patients** |  |  |  |  |  |
| COL6 FP0001 | COL6A2: het. c.2329 T>C<br>COL6A3: het. c.403 G>A | UCMD | Primary | 8.6 | Sant Joan de Déu |
| COL6 FP0016 | COL6A1: het. c.877 G>A | BM | Primary | 2.0 | Sant Joan de Déu |
| COL6 FP0016ed | CR RNA2* | BM | Primary | 3.2 | Sant Joan de Déu |
| COL6 FP0052 | COL6A2: hom. c.1970-9 G>A | UCMD | Primary | 1.9 | Sant Joan de Déu |
| COL6 FP0024 | COL6A1: het. c.1056+1 G>A | BM | Primary | 2.6 | Sant Joan de Déu |
| COL6 FP0005 | COL6A2: het. c.1970-9 G>A<br>COL6A2: het. c.115+2 T>C | UCMD | Primary | 1.5 | Sant Joan de Déu |
| *CR RNA2 - FP0016 COL6 cells edited with CRISPR/Cas9 RNA guide 2 (crRNA2) by Dr. López-Márquez in Sant Joan de Déu<sup>16</sup>; |  |  |  |  |  |

### Cell culture

Fibroblast were grown in Dulbecco’s Modified Eagle Medium (DMEM) supplemented with 20% Foetal Bovine Serum (FBS), 2% GlutaMax, and 1% Penicillin-Streptomycin-Glutamine (all from Gibco, Waltham, MA, US) and tested for mycoplasma contamination using Venor®GeM Classic kit (Minerva BioLabs, Berlin, Germany). Cells were cultured at 37°C in 5% CO_2_.

### Immunocytochemistry procedure

Skin fibroblasts were seeded at a density of 1×10^4^-2×10^4^ cells per well on 24-well plates containing glass coverslips that had been pre-coated with collagen I (Sigma-Aldrich, St. Louis, MO, US) in accordance with the manufacturer’s instructions. The growth medium was then replaced with medium enriched with ascorbic acid (L-ascorbic acid phosphate 2-magnesium, Sigma-Aldrich, St. Louis, MO, US) according to the experimental design. Following the designated incubation time in ascorbic acid, cells were rinsed with PBS (Gibco, Waltham, MA, US) and fixed in 4% paraformaldehyde (Sigma-Aldrich, St. Louis, MO, US) for 10 minutes, followed by four 5-minute-washes in PBS.

For visualising both intracellular and extracellular collagen VI, cells in some wells were permeabilised by washing them four times with 0.1% Triton X-100 (Sigma-Aldrich, St. Louis, MO, US) in PBS for 5 minutes. To observe only the extracellular collagen VI, those wells were washed four times with PBS alone. Subsequently, all samples were then blocked with Intercept® (PBS) blocking buffer (LI-COR Biosciences, Lincoln, NE, US) for 2 hours and immunolabelled with anti-collagen VI antibody (Merck Millipore, Burlington, MA, US, MAB1944, 1:2500) diluted in blocking buffer for 1 hour. After being washed four times in PBS, cells were blocked for 5 min with blocking buffer, followed by a 1 h incubation with an appropriate secondary antibody (1:500 in blocking buffer), while being protected from light.

The cells were then subjected to a 10-minuted wash with 0.05% Tween-20 in PBS, incubated in Hoechst 33342 solution (Thermo Fisher Scientific, Waltham, MA, US) diluted 1:1000 in PBS for 5 minutes, washed three times with 0.05% Tween-20 (Sigma-Aldrich, St. Louis, MO, US) in PBS, and once in PBS. Finally, slides were mounted using ProLong antifade mounting medium (Thermo Fisher Scientific, Waltham, MA, US) and kept at 4°C. Images were captured using a Leica DMIL-LED epifluorescence microscope with a 10x objective and LAS X software (Leica Microsystems, Wetzlar, Germany).

### Flow cytometry

Assays were performed according to the latest version of the protocol proposed by Kim et al. ^12^, with some modifications. Skin fibroblasts were seeded at a density of 1×10^6^-2×10^6^ cells in a 175 cm^2^ tissue culture flask. The growth medium was replaced with medium containing ascorbic acid 48 and 96 hours after seeding, and the cells were further incubated for 72h. Cells were then washed twice in PBS (Mg^2+^ and Ca^2+^ free, Gibco, Waltham, MA, US) and once in Cell Dissociation Buffer (enzyme free, PBS, Gibco, Waltham, MA, US). Cells were harvested with the Cell Dissociation Buffer, counted to obtain 0.5×10^6^cells/tube and fixed with 2% paraformaldehyde for 10 minutes on ice. The cells were then washed twice with PBS and centrifuged at 4°C for 5 minutes at 500g. Pellets were re-suspended and incubated with either a monoclonal primary antibody against collagen type VI (Merck Millipore, Burlington, MA, US, MAB1944, 1:250) or with a monoclonal primary antibody against fibroblast surface protein (Sigma-Aldrich, St. Louis, MO, US, F4771, 1:20) in PBS/0.1% FBS or PBS/0.05% FBS and Tween 20 (for permeabilisation) for 30 min on ice. A negative control was set up incubating cells without the primary antibody on ice. Cells were then washed with PBS/0.1% FBS with or without Tween-20, and then centrifuged at 500 g for 5 minutes at 4°C. Secondary antibodies (Alexa Fluor 488 goat anti-mouse IgM and Alexa Fluor donkey anti-mouse IgG1, Invitrogen Life Technologies, Carlsbad, CA, US) were diluted (1:500) with either PBS/0.1% FBS or PBS/0.05% FBS with Tween 20 and incubated for 20 min on ice. Subsequently, the cells were washed with PBS, centrifuged at 500g for 5 min at 4°C, resuspended in PBS, and analysed using a MACSQuant® X flow cytometer (MACSQuant®: Miltenyi Biotec, Cologne, Germany).

### In Cell/On Cell Western Assay (collablot)

Skin fibroblasts were seeded into collagen I-coated 96-well plates (Thermo Fisher Scientific, Waltham, MA, US) at a density designed to achieve roughly 1.6×10^4^ cells per well after 72 hours (refer to Figure 2 for procedure and plate setup in). Subsequently, the growth medium was replaced with L-ascorbic acid phosphate 2-magnesium (Sigma-Aldrich, St. Louis, MO, US) according to the experimental design. After the designated incubation period in the L-ascorbic acid phosphate 2-magnesium medium, cells were fixed in 4% paraformaldehyde for 10 minutes and washed four times for 5 minutes in PBS. Samples were then incubated with DNA stain DRAQ5 (Thermo Fisher Scientific, Waltham, MA, US) diluted 1:1000 in PBS for 1 hour, after which the plates were scanned using an Odyssey M imaging system (Odyssey M®: LI-COR Biosciences, Lincoln, NE, US). The cells were subsequently permeabilised and blocked as per immunocytochemistry sample preparation protocols and incubated overnight at 4°C with either an anti-collagen VI primary antibody or a Collagen Hybridising Peptide (CHP, 3Helix, Utah, US) in blocking buffer. As specified in the manufacturer’s indications, CHP should be heated at 80° for 10 minutes and immediately placed on ice for another 10 minutes prior to use. After incubation, cells were washed four times in PBS and incubated with the appropriate secondary antibodies (LI-COR Biosciences, Lincoln, NE, US) diluted 1:800 in blocking buffer for 1 hour. Cells were then washed three times in 0.05% Tween-20 in PBS, followed by two washes in 200µl PBS. The plates were scanned using an Odyssey M plate reader and data were analysed using Empiria Studio 2.3 software (LI-COR Biosciences, Lincoln, NE, US).

**Figure 2.**
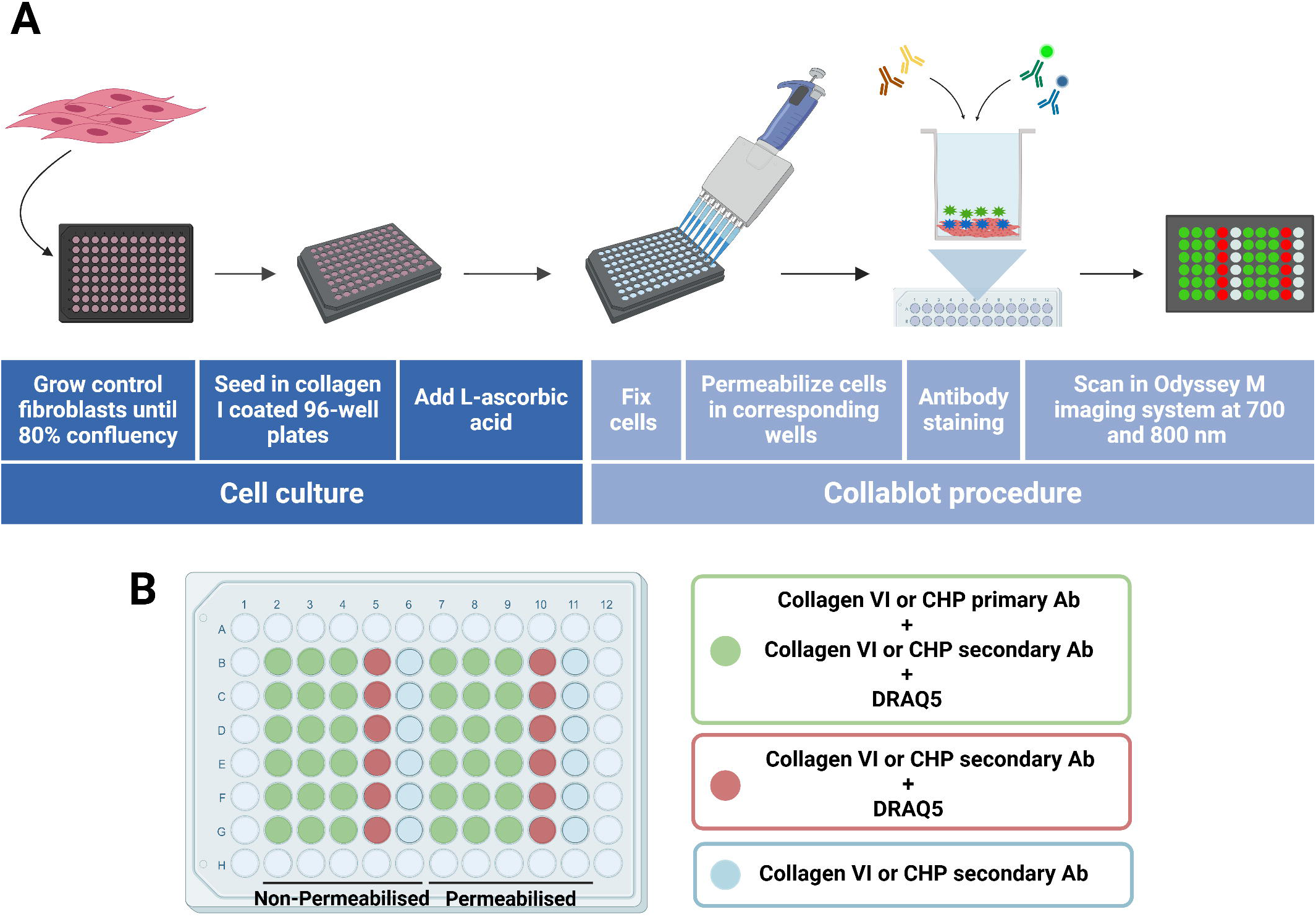
**A)** Illustration of collablot technique workflow. The first part of the workflow consists in cell culture and collagen secretion stimulation. The second is the collablot, where cells are fixed and incubated with a primary antibody, followed by its secondary antibody and then collagen or disorganisation is quantified. **B)** Set-up of the plates. Plates are divided in permeabilised and non-permeabilised. Wells in red indicate staining with DRAQ5 and wells in green staining with collagen VI antibody or CHP.

To ensure accurate quantification of both molecules, the analysis of CHP and collagen VI is conducted on separate wells. This approach is necessary due to the intercalating nature of CHP peptides, which may be hindered by antibodies binding to collagen. Wells stained to identify either collagen VI or CHP, which also include DRAQ5 for normalisation, are shown in green, wells in red indicate staining with DRAQ5 and the secondaries antibodies and wells in blue indicate no primary antibody control and no DRAQ5 (Figure 2). Water is added to the outer wells of all plates to prevent evaporation from the inner wells. The plate is divided such that columns 2-6 are non-permeabilised in order to analyse extracellular collagen while columns 7-11 are permeabilised in order to analyse total collagen. All readings were taken from the same plate utilising an Odyssey scanner at different wavelengths: 800CW for collagen VI, 520CW for CHP and 700CW for DRAQ 5.

The data collected via the Odyssey scanner were analysed as follows: first, the average of the readings from the corresponding no-primary background control wells was subtracted from each sample value. Following that, this background-corrected collagen VI or CHP data was normalised by dividing it by the DRAQ5 value of the same well. ((Value of a well in the 800CW or 520CW channel minus average nonprimary control wells)/value of the well in the 700 CW channel).

### Statistical analyses

Data were analysed using GraphPad Prism 9 (Graph-Pad Software Inc., La Jolla, CA, US). Student’s t-test were applied. Data are presented as mean +/- SEM. Differences were reported as significant as P < 0.05 (*), P < 0.01 (**) and P < 0.001 (***).

## Results

### Collablot optimization

#### Seeding density/Duplication times

Having a thorough understanding of cell cultures is essential when implementing any ICW technique, primarily due to the variability in doubling times amongst primary cultures. Table 1 summarises the doubling times of the cultures used in this study. When using cultures from different patients, adjustments must be made to account for the differences in doubling times, such as by varying the number of cells seeded per well.

To accurately quantify the DRAQ5 signal (700CW), used to normalise each well for cell density, it must fall within a lineal range. To stablish the number of cells needed to be within this range several cell dilutions were conducted. In these experiments, a control culture which displays intermediate growth characteristics, CTRL FP0821 (see supplementary Figure 1) was used. After studying this and other cultures, it was established that an initial seeding density of 6500 cells/well was appropriate for intermediate growth cultures (CTRL FP0821, CTRL FP, CTRL FP0010, CTRL FP0017, COLVI FP0016edit and COLVI FP0024), 5300 cells/well for high growth cultures (COLVI FP0016, COLVI FP0052, COLVI FP0005), and 12000 cells/well for low growth cultures (COLVI FP0001).

#### Selection of primary antibodies and collagen hybridising peptide

The basis of ICW and OCW assays are immunocytochemistry techniques, and a good selection of antibodies is necessary. The secondary antibodies and DRAQ5, were used at the concentrations suggested by each manufacturer. To select the most appropriate collagen VI primary antibody, a panel, consisting of three different antibodies previously validated for immunofluorescence (see supplementary Table 1) was evaluated. The antibodies compared were an anti-human collagen VI polyclonal antibody produced in rabbit (Abcam, Cambridge, UK: ab6588), and two anti-human collagen VI antibodies produced in mice (Merck Millipore, Burlington, MA, US: MAB3303 and MAB1944). ^6,9,11,12,17–26^

The ICW/OCW methodology detects native protein, as samples are not lysed nor denaturalised. Therefore, it is advisable to evaluate the antibodies through a standard immunocytochemistry experiment. The CTRL FP0821 cell line was probed with the three primary antibodies (Supplementary Table 1) and their respective secondary antibodies, Alexa Fluor 488 anti-rabbit IgG (Fisher Scientific, Hampton, NH, US) for ab6588, and Alexa Fluor 488 anti-mouse IgG (Fisher Scientific, Hampton, NH, US) for MAB3303 and MAB1944. As illustrated in Supplementary Figure 2, while the anti-rabbit secondary antibody showed a non-specific signal in the absence of the primary antibody, the anti-mouse secondary antibody shows no such non-specific signal, confirming its suitability for use. The polyclonal antibody ab6588 displayed minimal signal, and the faint signal detected could be attributed to the non-specific signal generated by the secondary antibody. In contrast, both monoclonal primary antibodies demonstrated a positive response, thus, warranting further evaluation (Figure 3A).

**Figure 3.**
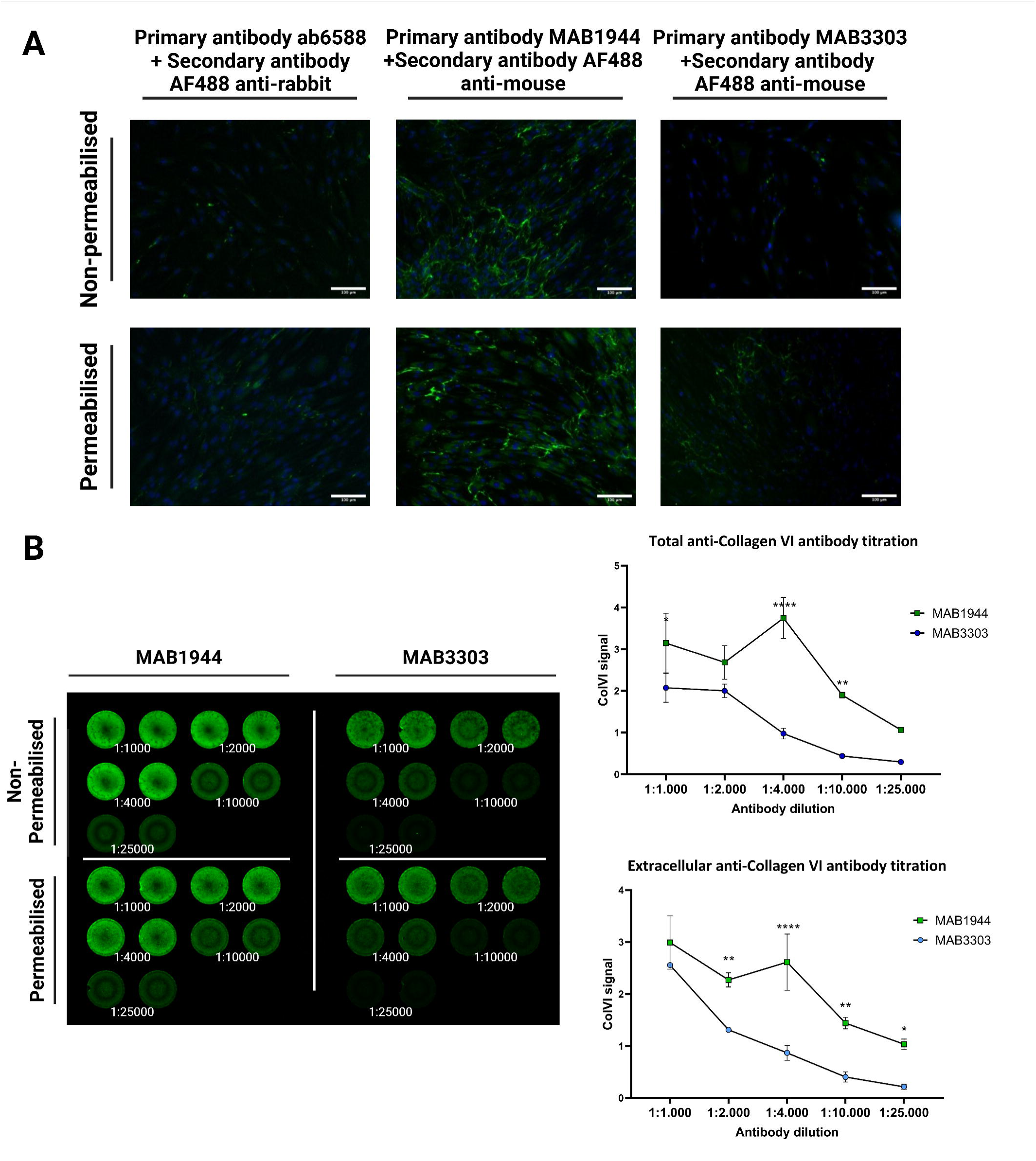
**A)** Immunocytochemical analysis of the three different primary antibodies for collagen VI (ab6588, MAB1944 and MAB3303; 1:2500) carried out using CTRL FP0821 line, permeabilised and non-permeabilised. A single replicate with two repeats. **B)** On-cell western analysis with two different primary antibodies for collagen VI (MAB1944 and MAB3303) titrations (1:1000, 1:2000, 1:4000. 1:10000 and 1:25000) carried out using CTRL FP0821 line, permeabilised (total collagen) and non-permeabilised (extracellular collagen). A single replicate with two repeats.

To ascertain the more suitable monoclonal antibody for this type of assay and its optimal titration, an ICW/OCW assay was conducted using the CTRL FP0821 cell line and various antibody dilutions (1:1000, 1:2000, 1;4000, 1:10000 and 1:250000). The MAB1944 antibody exhibited a higher signal compared to the MAB3303 antibody at all tested concentrations, corroborating the results obtained in the immunohistochemistry assay (Figure 3B). Furthermore, including both permeabilised and non-permeabilised cells, the greatest signal-to-background ratio was achieved with the 1:4000 dilution. Consequently, the primary antibody MAB1944 at a dilution of 1:4000 was selected.

In addition to the amount of collagen produced and its distribution inside or outside cells, the level of disorganization in the collagen filaments might also give valuable insights about the cultures. Therefore, a peptide capable of intercalating among loosely bound collagen fibres (CHP) was assessed (Figure 4A). To assess whether the CHP colocalised with collagen VI, an immunocytochemical analysis was conducted (Figure 4B). To determine the optimal titration of CHP to use, another ICW/OCW assay was conducted on the same control line, using different CHP dilutions (1:5, 1:10 and 1:20) (Figure 4C) selected taking into a count the manufacturer’s indications, where CHP titration should be between 5 and 30 µM. Considering both permeabilised and non-permeabilised cells, the highest signal-to-background ratio was achieved with the 1:10 dilution. Therefore, the CHP dilution of 1:10 was selected.

**Figure 4.**
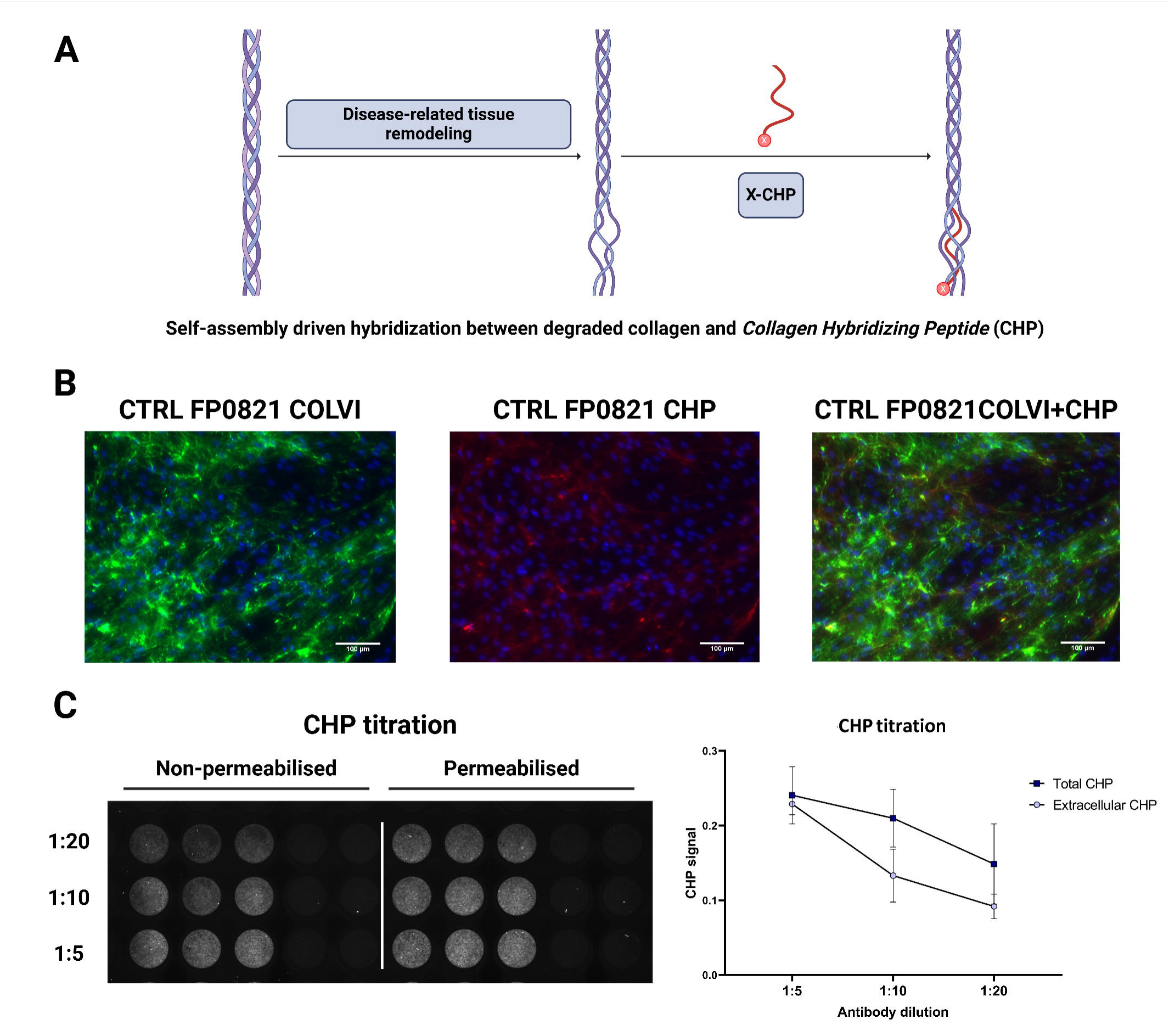
**A)** Mechanism of action of the Collagen Hybridising Peptide (CHP). **B)** Immunocytochemical analysis of the colocalisation of the CHP (1:5) and the collagen VI (1:2500) carried out using non-permeabilised CTRL FP0821 line. A single replicate with two repeats. **C)** On-cell western analysis of CHP titrations (1:5, 1:10 and 1:20) carried out using CTRL FP0821 line, permeabilised and non-permeabilised. A single replicate with three repeats.

#### Optimization of L-ascorbic acid concentration for collagen VI secretion

Ascorbic acid is essential for collagen VI synthesis and promotes the secretion of this protein onto the extracellular medium. An immunocytochemical assay (Supplementary Figure 3) was performed to determine the best incubation time and concentration of ascorbic acid required to detect variations in collagen VI synthesis and secretion. This experiment confirmed that both ascorbic acid concentration and incubation time are crucial factors affecting collagen secretion: the higher these parameters, the greater the secretion. We were able to quantify this observation using the ICW/OCW (collablot) assay (Figure 5): with this technique it was possible to visualise not only the optimal combination of a concentration of ascorbic acid of 50 μg/μL with a 48-hour incubation, but also demonstrated the collablot’s ability to detect various concentrations of collagen both extracellularly and intracellularly.

**Figure 5.**
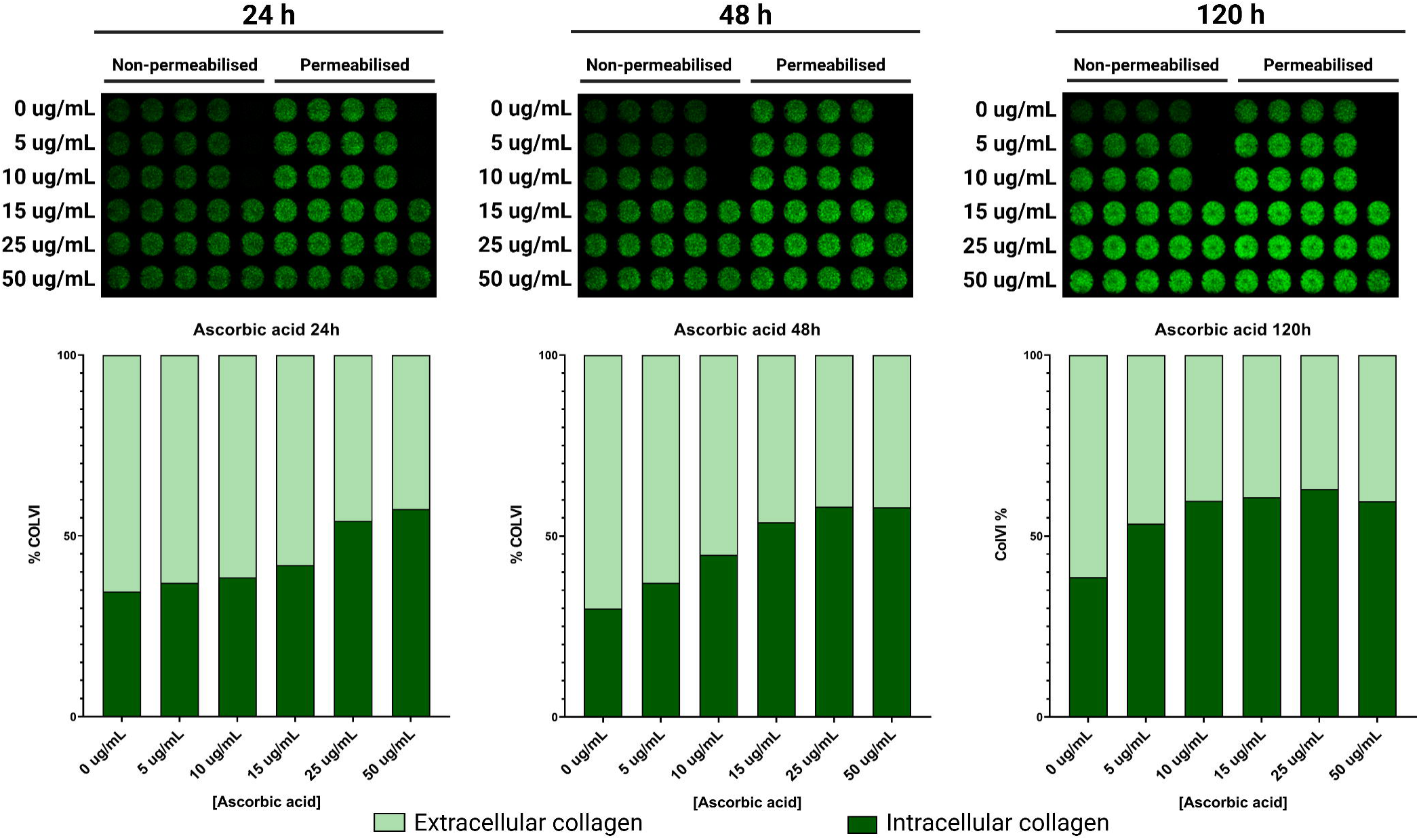
On-cell western analysis with the same parameters than the ICC and the quantitative results of both intracellular and extracellular collagen VI expressed due to the effect of the ascorbic acid. A single replicate with four repeats.

### Collablot of control and patient cultures

Cultures derived from control and patient biopsies were analysed using our optimised collablot method (Figure 6). The immunocytochemical assay (Figure 6A) provides qualitative data, while the collablot quantifies extracellular and total collagen (Figure 6B). The quantification of collagen with the collablot mirrored the immunohistochemistry results: while a trend for lower collagen content and a larger percentage of intracellular collagen can be detected, it is not possible to statistically distinguish patients from control groups, due to the heterogeneity present amongst patients and highlighting the need for an additional analysis.

**Figure 6.**
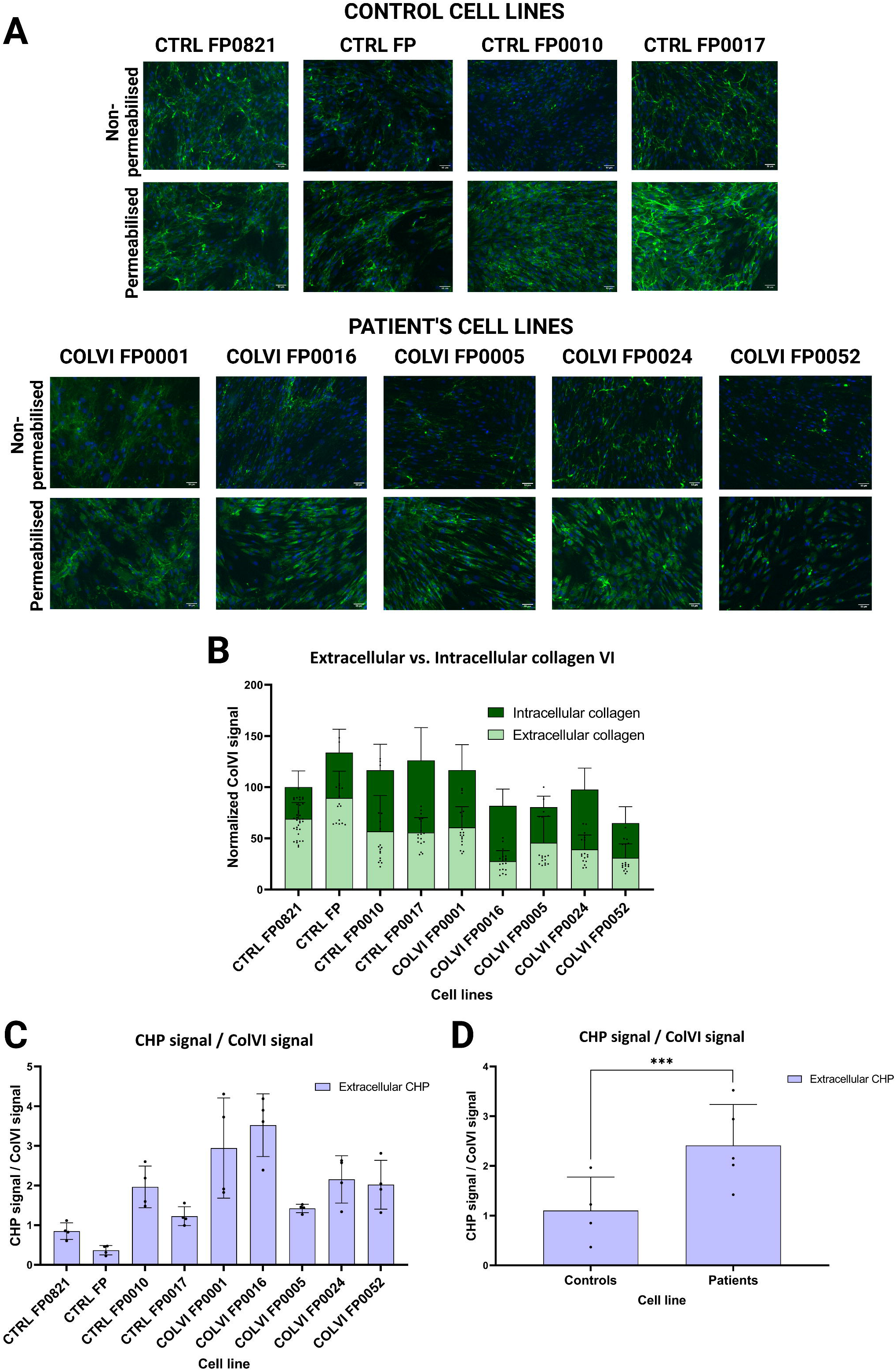
**A)** Immunocytochemical results of collagen VI for each cell line. Two replicates with two repeats per replicate. **B)** On-cell Western results comparing total and extracellular collagen from all cell lines. Six replicates with six repeats per replicate. **C)** Extracellular CHP signal normalised by extracellular ColVI signal obtained from on-cell western experiments of all cell lines. Four replicates with three repeats per replicate. **D)** Extracellular CHP signal normalised by extracellular ColVI signal of controls (CTRL FP0821, CTRL FP, CTRL FP0010 and CTRL FP0017) and patients (COLVI FP0001, COLVI FP0016, COLVI FP0005, COLVI FP0052 and COLVI FP0024) cell lines. Same replicates and repeats as in Figure 6C.

The quantification by collablot of the CHP peptide signal in these cultures is indicative of the levels of organization of the extracellular collagen matrix and showed that the levels of CHP are diverse among the different cell lines tested. As this signal is also dependent on the total amount of collagen, it was shown relative to the total collagen amount present in each culture (Figure 6C). When the data was displayed in this way, control cultures presented a lower CHP/ColVI ratio when compared to patient cultures (Figure 6D), corresponding with the amount of peptide being able to intercalate within the fibres due to disorganisation.

### Evaluation of collagen VI therapeutic restoration by collablot

As one of the objectives of this study is to develop a method that could be used to screen new treatments, we used it to evaluate an isogenic cell culture pair: a “rescued” culture was created after gene edition of a patient’s fibroblast culture and these cultures were kindly provided by HSJD and analysed in Biobizkaia. The cultures had been characterised in the original publication, and we confirmed that characterisation by subjecting the cultures to analysis by immunocytochemistry (Figure 7A), collagen collablot (Figure 7B), flow cytometry (Figure 7C) and, and CHP analysis by collablot (Figure 7D).

**Figure 7.**
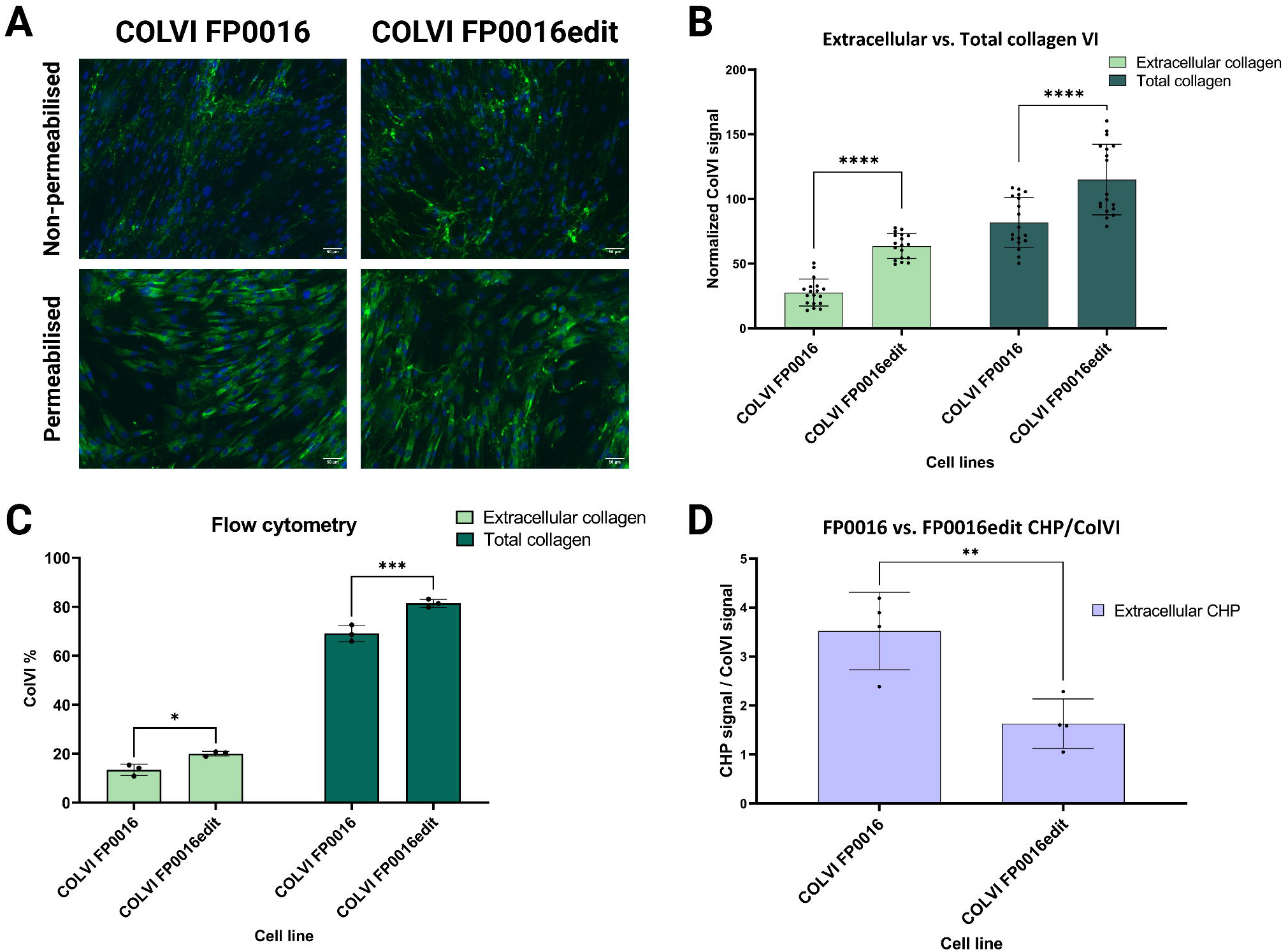
**A)** Immunocytochemical results for COLVI FP0016 and its edited line, COLVI FP0016edit, extracellular and total collagen. **B)** On-cell western results of COLVI FP0016 cell line and its edited line, COLVI FP0016edit. Six replicates with three repeats per replicate. **C)** Flow cytometry results of COLVI FP0016 cell line and its edited line, COLVI FP0016edit. A single replicate with three repeats. **D)** Extracellular CHP signal normalised by extracellular ColVI signal of COLVI FP0016 cell line and its edited line, COLVI FP0016edit. Four replicates with three repeats per replicate.

It was possible to corroborate the restoration of collagen expression described in the original publication by immunohistochemistry, flow cytometry and collagen collablot. With this technique, it was confirmed that both total and extracellular collagen signal were higher in the edited cultures. Additionally, CHP quantification by collablot confirmed a very significant decrease in the extracellular fraction, corresponding to a lower amount of CHP bound to denatured collagen strands suggesting that after the edition of the COLVI FP0016 cell line, the collagen matrix is more organised and structured.

## Discussion

The diagnosis of myopathies associated with collagen deficiency requires, in addition to clinical assessment, confirmation of collagen deficiency is a complementary assay to genetic analysis of the *COL6A1*, *COL6A2* and *COL6A3* genes. This is a problem because most diagnostic techniques are not quantitative and, those that are, do not accurately distinguish patients with the milder versions of the disease, as their collagen levels are indistinguishable from controls. The only quantitative method capable of distinguishing BM from UCMD reported so far is flow cytometry. However, this is an indirect quantification, as it measures the number of collagen-expressing cells, not the amount of collagen excreted^12^.

The structural and organisational consequences of *COL6* mutations depend on their location. Those at the N-terminal of the triple helix domain always have a negative impact on the assembly of tetramers and microfibrils. However, for mutations located outside the triple helix domain, the disorganisation of the collagen network depends on whether they prevent incorporation into monomers and are then targeted for proteasomal degradation, or whether they are incorporated into monomers, dimers and/or tetramers that disrupt the formation and/or organisation of microfibrils^27,28^.

We have now developed a method that we think may contribute to a faster development of new treatments for neuromuscular disorders, as it allows a more accurate quantification of collagen VI with the use of a limited number of cells. Unfortunately, collagen quantification alone would not suffice as a diagnostic tool because there are patients who express equal or greater amounts of both extracellular and intracellular collagen compared to some of the controls. This suggests that the disease is not solely determined by the quantity of collagen, and the organization of collagen also plays a significant role in the disease. Some studies^29–32^ have demonstrated the ability of the collagen hybridising peptide (CHP) to bind unfolded collagen chains and thus show damage at the molecular level. The incorporation of this molecule into our technique has demonstrated a large disruption of the collagen network in patients relative to the control cohort. While CHP quantification is not sufficient for a definitive diagnosis of COL6-RD, in combination with the quantification of collagen expression provides a valuable insight into the tissue damage present in the patient under study.

Even though collablot method is a robust tool that provides the quantification of both collagen VI and the organization levels of the extracellular collagen; is it likely not enough for the diagnosis of this disorder. The myopathies caused by collagen deficiencies are complex conditions and there are several other factors that could influence its diagnosis. One key factor is the penetrance of the mutations, the large number of variants of uncertain meaning and the frequent mosaicism found in this type of disease^1^.

Nevertheless, the collablot method is a quantifiable and useful method for the evaluation of novel therapies, as demonstrated in this manuscript with the characterisation of a cell culture isogenic pair one including a mutation in the *COL6A1* gene, and an edited version created by CRISPR/Cas9^16^. The method we describe easily showed how collagen VI levels were increased in the edited cell line compared to the original cell line from the patient. And, more interestingly, the analysis of CHP levels showed a very marked improvement in the organisation of that collagen VI in the extracellular matrix, essential point for this disease.

Our results show that collablots could be a fast and suitable technique for the quantification of both collagen VI and its disorganisation in skin-derived fibroblasts. Although this quantification is not sufficient for disease diagnosis on its own, it can provide new insights available to clinicians and it is an optimal method for assessing the impact of therapies on collagen expression. We expect that this technique will be useful for the neuromuscular community.

## Acknowledgements

We strongly acknowledge Fundación Noelia for supporting this work and for its enormous effort in securing funding, their trust in our work, and extraordinary cooperation.

We acknowledge the use of cell cultures provided by Dr López de Munain, Biogipuzkoa Health Research Institute (Donostia-San Sebastián, Spain), and the Biobank (BHISJDI) of the Hospital Sant Joan de Déu (HSJD) (Barcelona, Spain).

## Data availability

All relevant data can be found within the article and its supplementary information. All materials and further information of this study is available upon request.

## Author contribution statement

Conceptualization, V.A.-G.; methodology, validation, formal analysis and investigation, resources, V.A.-G.; writing-original draft preparation, M.S; N. O-B.; writing-review and editing, A. L.-M.; C. J.-M.; A. L.-M.; M.S.; I.G.-M.;-V.A.-G.; visualisation, supervision, A. L.-M. M.S.; I.G.-M.; V.A.-G; project administration and funding acquisition, V.A.-G.

## Abbreviations

COL6-RD: Collagen VI-related dystrophies
BM: Bethlem myopathy
UCMD: Ulrich congenital muscular dystrophy
ECM: Extracellular matrix
IHC: Immunohistochemistry
ICC: Immunocytochemistry
WB: Western Blot
OCW: On-cell western
ICW: In-cell western
FC: Flow Cytometry
CHP: Collagen Hybridising peptide

---

***Supplementary Figure 1.*** Lineal range of data from collablots. Serial dilutions (3000-10000 cells per well) of non-permeabilised and permeabilised CTRL FP0821 fibroblasts analysed in a collablot. A single replicate with three repeats.

***Supplementary Table 1.*** Anti-collagen VI primary antibodies tested in this project and its molecular localisation in the three different alpha chains of collagen VI (α1(VI), α2(VI) and α3(VI)).

***Supplementary Figure 2.*** Immunocytochemical analysis of the two different secondary antibodies (Alexa Fluor 488 anti-rabbit IgG and Alexa Fluor 488 anti-mouse IgG) for collagen VI primary antibodies (ab6588, MAB1944 and MAB3303; 1:2500) carried out using CTRL FP0821 line, permeabilised and non-permeabilised. A single replicate with two repeats.

***Supplementary Figure 3.*** Immunocytochemical analysis of permeabilised and non-permeabilised CTRL FP0821 line at 24, 48 and 120 hours with different concentrations of ascorbic acid (0, 5, 10, 15, 25 and 50 µg/mL) and a negative control for each time. (NP-Non permeabilised; P-Permeabilised). A single replicate with two repeats.

