## Supplementary figures for "Collablots: Quantification of collagen VI levels and its structural disorganisation in cell cultures from patients with collagen VI-related dystrophies"

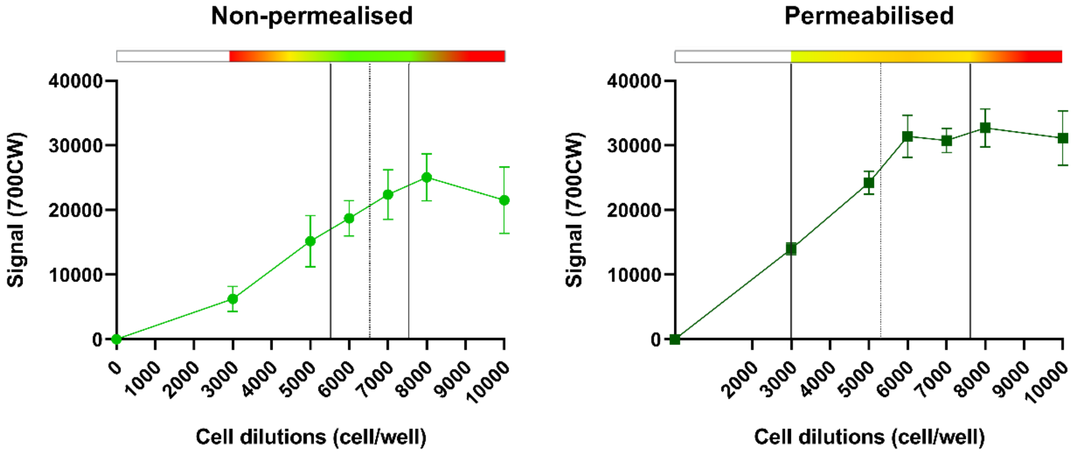


***Supplementary Figure 1***. Lineal range of data from collablots. Serial dilutions (3000-10000 cells per well) of non-permeabilised and permeabilised CTRL FP0821 fibroblasts analysed in a collablot. A single replicate with three repeats.

***Supplementary Table 1.*** Anti-collagen VI primary antibodies tested in this project and its molecular localisation in the three different alpha chains of collagen VI (α1(VI), α2(VI) and α3(VI)).

| Antibody | Supplier | Cat number | Isotype and host | Specificity | References |
|---|---|---|---|---|---|
| **Anti-collagen VI antibody** | Abcam | ab6588 | Rabbit IgG | Non-helicoidal region of α1(VI) | **IHC** [11, 17, 18]<br>**ICC** [19, 20] |
| **Anti-collagen VI antibody, clone VI-26** | Millipore | MAB3303 | Mouse IgG1 | Triple helix of α1(VI), α2(VI) and α3(γ) | **IHC** [21, 22]<br>**ICC** [23, 24]<br>**WB** [9.] |
| **Anti-collagen VI antibody, clone 3C4** | Millipore | MAB1944 | Mouse IgG1 | Non-helicoidal region of α3(VI) | **IHC** [21, 22]<br>**ICC** [6, 25, 26.]<br>**WB** [9]<br>**FC** [12] |

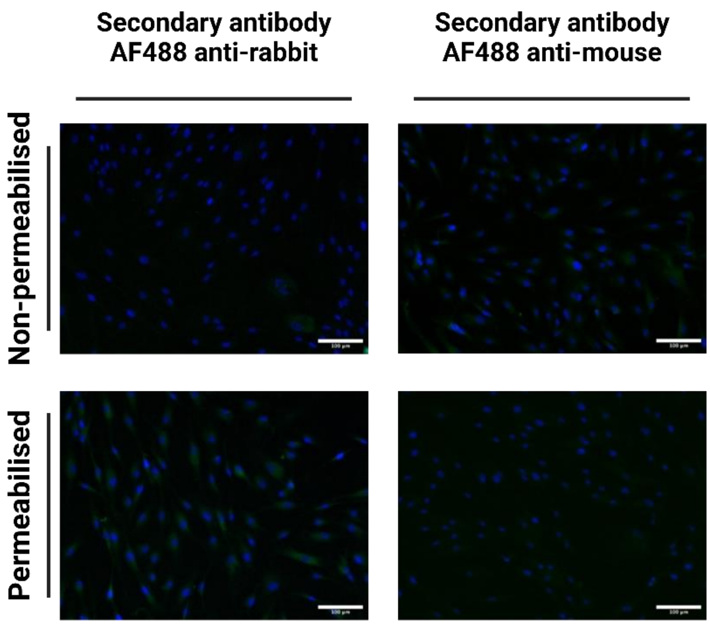


***Supplementary Figure 2.*** Immunocytochemical analysis of the two different secondary antibodies (Alexa Fluor 488 anti-rabbit IgG and Alexa Fluor 488 anti-mouse IgG) for collagen VI primary antibodies (ab6588, MAB1944 and MAB3303; 1:2500) carried out using CTRL FP0821 line, permeabilised and non-permeabilised. A single replicate with two repeats.

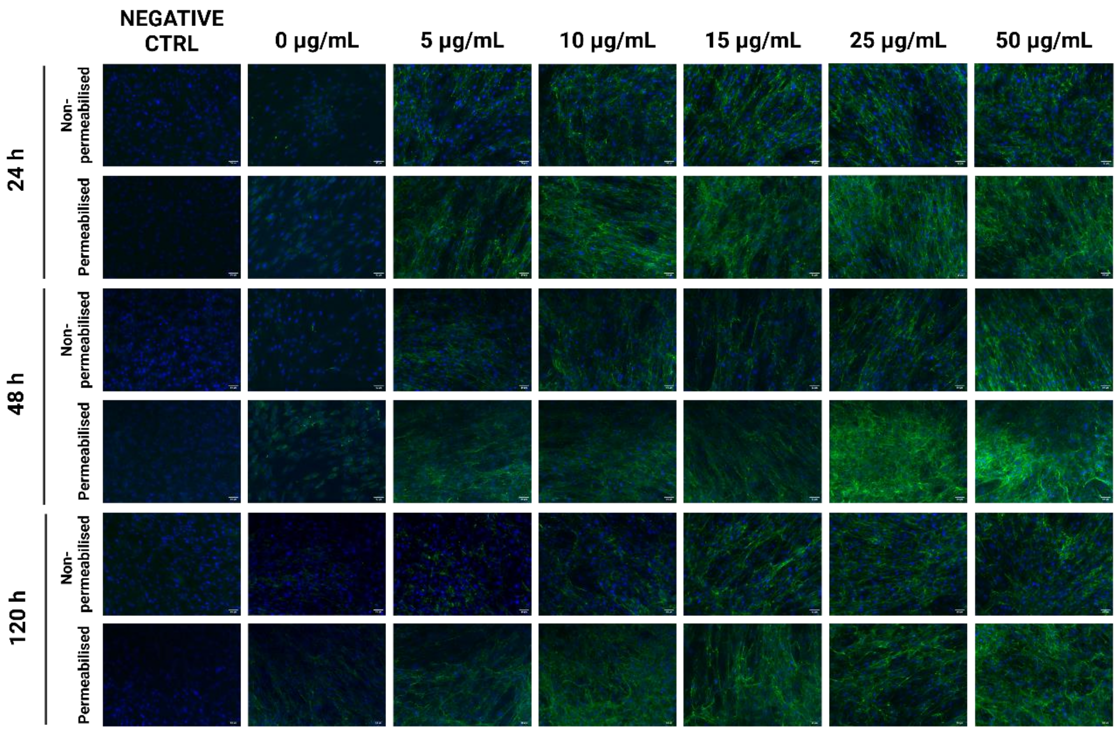


***Supplementary Figure 3.*** Immunocytochemical analysis of permeabilised and non-permeabilised CTRL FP0821 line at 24, 48 and 120 hours with different concentrations of ascorbic acid (0, 5, 10, 15, 25 and 50 µg/mL) and a negative control for each time. (NP-Non permeabilised; P-Permeabilised). A single replicate with two repeats.
